# Modeling consensus and distance to delivering indoor residual spray (IRS) implementation strategies to control malaria transmission

**DOI:** 10.1101/2020.05.07.082982

**Authors:** Sadie Ryan, Anne C. Martin, Bhavneet Walia, Anna Winters, David A Larsen

## Abstract

**Background:** Indoor residual spraying (IRS) is an effective method to control malaria-transmitting *Anopheles* mosquitoes and often complements insecticide-treated mosquito nets, the predominant malaria vector control intervention. With insufficient funds to cover every household, malaria control programs must balance the malaria risk to a particular human community against the financial cost of spraying that community. This study creates a framework for modeling the distance to households for targeting IRS implementation, and applies it to potential risk prioritization strategies in four provinces (Luapula, Muchinga, Eastern, and Northern) in Zambia.

**Methods:** We used optimal network models to assess the travel distance of routes between operations bases and human communities identified through remote sensing. We compared network travel distances to Euclidean distances, to demonstrate the importance of accounting for road route costs. We then compared the distance to reaching communities for different risk prioritization strategies assuming sufficient funds to spray 50% of households, using four underlying malarial risk maps: a) predicted *Plasmodium falciparum* parasite rate in 2-10 year olds (*Pf*PR), or b) predicted probability of the presence of each of three main malaria transmitting anopheline vectors (*An. arabiensis, An. funestus, An. gambiae*)

**Results:** The estimated one-way network route distance to reach communities to deliver IRS ranged from 0.05 – 115.69 Km. Euclidean distance over and under-estimated these routes by −101.21 – 41.79 Km ***per trip***, as compared to the network route method. There was little overlap between risk map prioritization strategies, both at a district-by-district scale, and across all four provinces. At both scales, agreement for inclusion or exclusion from IRS across all four prioritization strategies occurred in less than 10% of houses. The distances to reaching prioritized communities were either lower, or not statistically different from non-prioritized communities, at both scales of strategy.

**Conclusion:** Variation in distance to targeted communities differed depending on risk prioritization strategy used, and higher risk prioritization did not necessarily translate into greater distances in reaching a human community. These findings from Zambia suggest that areas with higher malaria burden may not necessarily be more remote than areas with lower malaria burden.

## Background

Indoor residual spray (IRS) is an effective method to control the *Anopheles* mosquitoes that transmit malaria [1]. The intervention has helped drive success in decreasing malaria transmission across sub-Saharan Africa [2,3]. IRS is often seen as complementary to the use of insecticide-treated mosquito nets (ITN), which is the predominant vector control intervention to prevent malaria transmission [3,4].

In contrast to ITNs, which in 2011 cost an estimated $2.20 per year of protection delivered, IRS was much more expensive [5]. Using the chemicals DDT, pyrethroids, deltamethrin, and lambdacyhalothrin, the cost of IRS was $6.70 per year of protection per household, with the cost of insecticide ranging from 29 – 81% of the total cost and minimal economies of scale [5]. With second generation ITNs and the drive for lower insecticides for IRS these prices have changed somewhat over the past 10 years. Pyrethroids have decreased in cost substantially over the past 25 years [6], but widespread insecticide resistance threatens the long term viability of using pyrethroids for malaria control [7–9]. Currently no alternative insecticides for ITNs are available for use at scale, although novel chemicals and dual-chemical ITNs are in various stages of development. Alternative insecticides for IRS are available but come at a greater cost than those of DDT, pyrethroids, deltamethrin, and lambdacyhalothrin. Indeed, IRS programs funded by the organizations such as the US President’s Malaria Initiative (PMI) have seen reductions in coverage due to the increasing cost of insecticide [10].

Often countries are faced with the challenge of insufficient funds to cover every household in malaria endemic areas, and as such are forced to determine which houses receive the intervention. Zambia’s National Malaria Elimination Centre (NMEC) has encountered this challenge; in at least some areas health facility malaria incidence is used to prioritize areas to receive IRS [11]. Other approaches have been used to prioritize which areas receive IRS, but whichever methodology is used, malaria programs must balance the malaria risk of a particular human community with the financial cost of spraying a particular human community.

The use of network modeling for optimizing the distribution of goods along road networks is commonly used for market analyses [12,13], and has been applied in the health arena to least-cost routing for hospital access [14–16] and delivery of vaccines [17,18]. Several authors have noted that a primary limitation to application of network modeling in the developing world is the availability of accurately mapped road networks [19,20]. This article creates a framework for modeling the cost of IRS implementation and applies that framework to potential intervention prioritization strategies in Zambia.

## Methods

### Study area

Zambia lies in southern Africa and has a range of malaria transmission intensity, from pre-elimination status in Southern and Lusaka provinces, to intense malaria transmission in Luapula Province. The modern history of indoor residual spray (IRS) in Zambia began in 2003 when the Government of the Republic of Zambia (GRZ) began spraying to complement the private sector’s IRS campaigns. Zambia’s approach is to support as many districts as possible with IRS; resources often do not allow for complete coverage of districts. This study focuses on four provinces in the eastern part of the country: Luapula, Northern, Muchinga, and Eastern provinces (Figure 1). Malaria indicator surveys estimate *Plasmodium falciparum* parasite prevalence rate to be > 25% and household ownership of at least one ITN > 50% in these areas [21].

**Figure 1.**
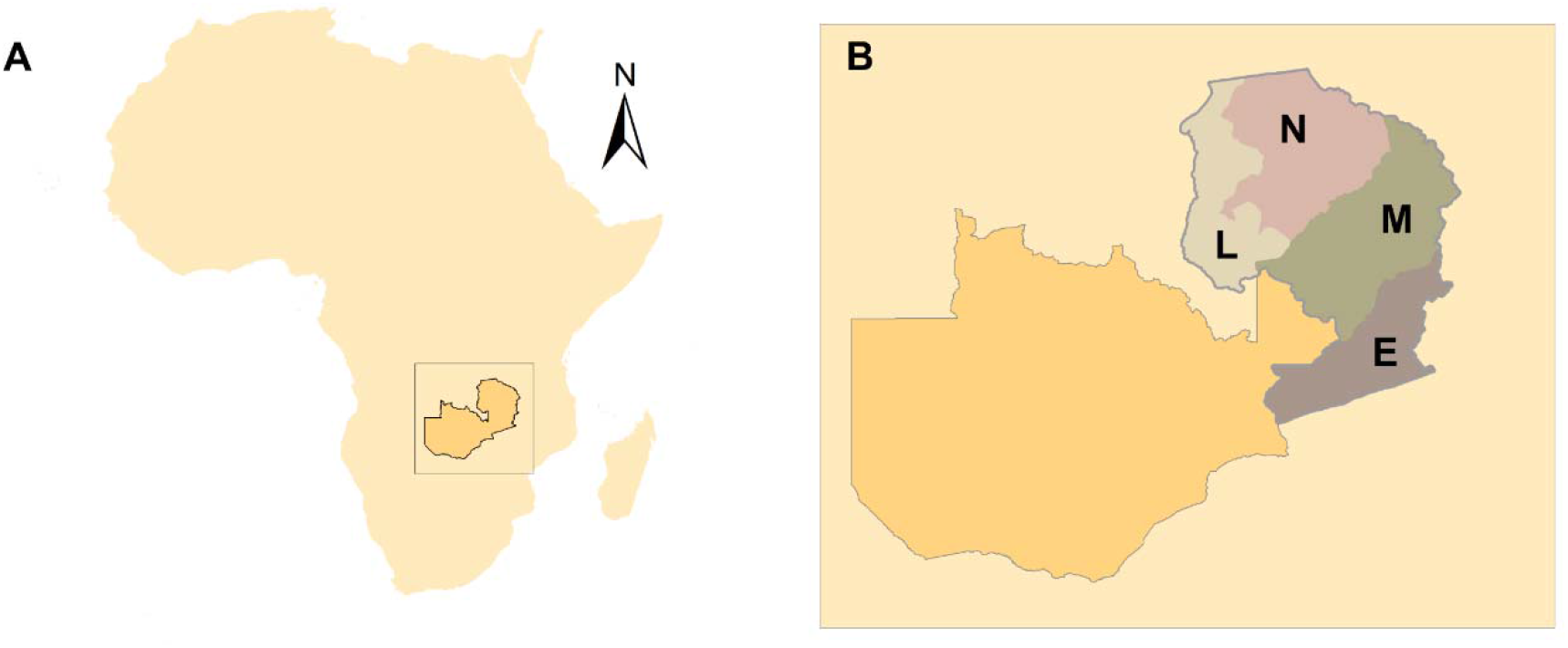
The location of A. Zambia in Africa, and B. the four provinces in this study L: Luapula, N: Northern, M: Muchinga, E: Eastern.

### Community data

Technicians in Lusaka, Zambia digitized structures in Eastern, Luapula, Muchinga, and Northern province visible in publicly available satellite imagery as part of the planning process of IRS campaigns in 2015 and 2016, as has been described elsewhere [22]. As part of the planning process of IRS campaigns, digitized structures were spatially aggregated into communities based upon distance between structures (< 50m), and then communities with fewer than 25 houses were deemed too small for IRS [11], and not included in the modeling. Operation bases were assumed to be located at city/village centers, which were taken as point locations from Google Map Maker [see next section for details], totaling 236 locations across the four provinces in this analysis.

### Road network data

Road network data were made available to this analysis via Google Map Maker. While Google Map Maker was an openly collaborative resource during the course of this study, it was retired in March, 2017 [23,24], and approved contributions merged with Google Maps. Google Map Maker data were chosen due to more complete road network mapping than OpenStreetMap (OSM) [25], or any other digital road map resource available at the time. For the four provinces, the mapmaker (MM) road file used in this study contained a total of 13,762,691.86 meters of roads, in 21,082 polyline segments (average segment length 653m). While MM road attributes describe primary and local roads separately, the absence of descriptors for local roads beyond identification meant that all roads in the network were treated equally.

### Network analysis

For each province, the road network was converted to a network database in ArcGIS 10.1’s Network Analysis toolbox. Impedence was set using meters. The city centers were added to the Arc project, and imported as ‘facilities’ for a Closest Facility analysis. The Closest Facility analysis is based on Dijkstra’s algorithm [26] to find the shortest path between two specified nodes on a network, thus providing optimal routing. This algorithm is ‘greedy’ in that it requires considerable computation; in essence, it calculates the distance between all nodes on a network, along all possible routes, and then for each node pair, reports the minimum of those distances. In ArcGIS 10.1’s Network Analysis toolbox, these routes are also saved as polyline files. In this analysis, the shortest route between each target centroid and its nearest city center was returned; a modification of Dijkstra full set.

To describe the communities, the centroid for each community served as a destination node on the network. As many of these were not on the road network, a tolerance of inclusion in the network of 5,000 m, both for the city center locations and for the community centroid locations, allowed for more communities in the analysis. Since the communities were not all within 5,000 m of the available road network, the network analysis was conducted on a reduced set of targets and city centers, as described in Additional Table 1. Network modeling was conducted in ArcGIS 10.1’s Network Analysis toolbox.

In addition to the network modeled route costing, because we set a tolerance of 5,000m, we added a Euclidean distance measure from each target centroid to the nearest point on the nearest road in the network, and calculated the total distance along the optimal route plus distance to the nearest road, to estimate true distance to the target centroid point (n=11,146).

To examine the impact of modeling distribution routes as optimal network routes, rather than simple Euclidean distance routes between communities and their nearest city centers, estimates of the two distances for each of the four provinces were compared.

### Establishing spatial malarial risk prioritization strategies

The distance framework was applied to two separate prioritization strategies based on underlying risk maps, namely: *Plasmodium falciparum* prevalence rate among kids aged 2-10 years old (*Pf*PR_2-10_) from the Malaria Atlas Project, estimated for 2010 [27]; and Malaria Atlas Project vector suitability for *Anopheles gambiae, Anopheles funestus*, and *Anopheles arabiensis* [28]. Malaria Atlas Project estimates of both *Pf*PR_2-10_ and vector suitability estimates were extracted from raster files to community polygons, and aggregated to average values, using the Raster package [29,30] in R version 3.3.2 [31].

### Analysis

Two separate modeling strategies were employed, the “within district prioritization” and the “across district prioritization”. For the first, half of all households *within* each district of the four provinces were targeted for IRS with prioritization based on one of the underlying risk maps.

Second, half of all households *across the four provinces* were targeted for IRS with the same underlying risk map prioritization. Prioritization was conducted by ranking communities in order of risk (high to low), and summing household numbers (counts within communities) from the top-ranked down, until 50% was exhausted. *t*-tests were used to determine differences by prioritization strategy and Kappa test scores were used to estimate the agreement between prioritization strategies. To generate a measure of overall agreement between the different prioritization strategies the arithmetic mean between pair statistics was taken [32]. Stata version 13.1 was used for these analyses.

## Results

### Mapping prioritized targets to a road network

The road network data used in this study represents only around a third of visible roads and tracks on the ground (Ryan, *unpublished*), and therefore should be considered a primary road network, rather than a full road network. Among the 18,448 communities created from the enumeration process, we were able to capture 11,146 (60.4%) to the incomplete road network. Of these communities, 3,136 had at least 25 houses and were included in further analysis. There was minimal difference in risk estimates between captured and non-captured communities based on *Pf*PR, *An. arabiensis*, and *An. funestus*, and captured communities had somewhat higher estimates of risk based on *An. gambiae* (Table 1). Captured communities were also on average larger than non-captured communities in terms of the number of houses in each community.

**Table 1.**
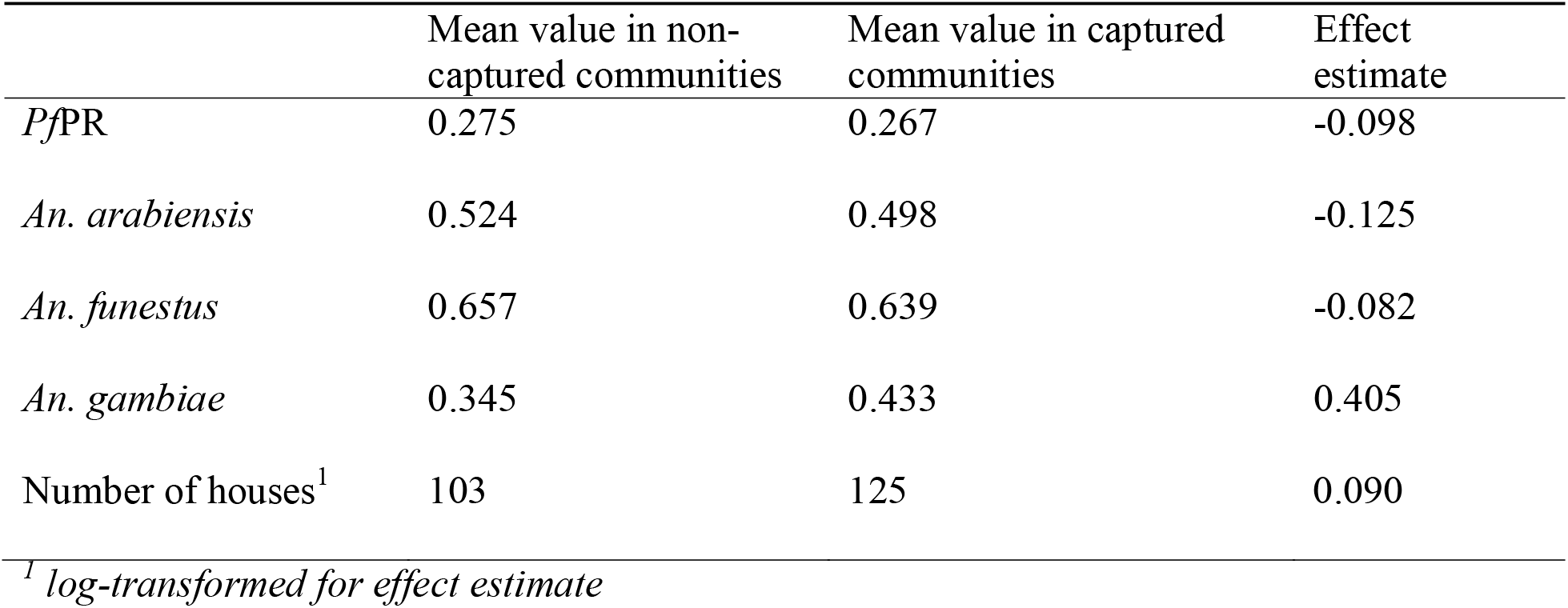
Differences in risk estimates (*Pf*PR, *An. arabiensis, An. funestus, An. gambiae*) between communities captured to road networks and those not captured to road networks.

|  | Mean value in non-captured communities | Mean value in captured communities | Effect estimate |
| --- | --- | --- | --- |
| *PfPR* | 0.275 | 0.267 | -0.098 |
| *An. arabiensis* | 0.524 | 0.498 | -0.125 |
| *An. funestus* | 0.657 | 0.639 | -0.082 |
| *An. gambiae* | 0.345 | 0.433 | 0.405 |
| Number of houses<sup>1</sup> | 103 | 125 | 0.090 |

<sup>1</sup> *log-transformed for effect estimate*

### Road Network versus Euclidean Routing

The differences between Euclidean (“straight-line”) and optimal road network route distances for each province are shown as histograms in Figure 2, A-D. These show that both over and underestimates arise under a simple Euclidean distance modeled view, generating an overall range of a maximum of 41.79 Km (overestimate) and a minimum of −101.21 (underestimate) of the distance of a one-way route to deliver spray to a community. While the overestimates range a substantial amount, overestimation at least errs in a conservative direction for operational planning. However, the underestimates, if part of multiple round-trips, could rapidly amount to large costs for local spray operations.

**Figure 2.**
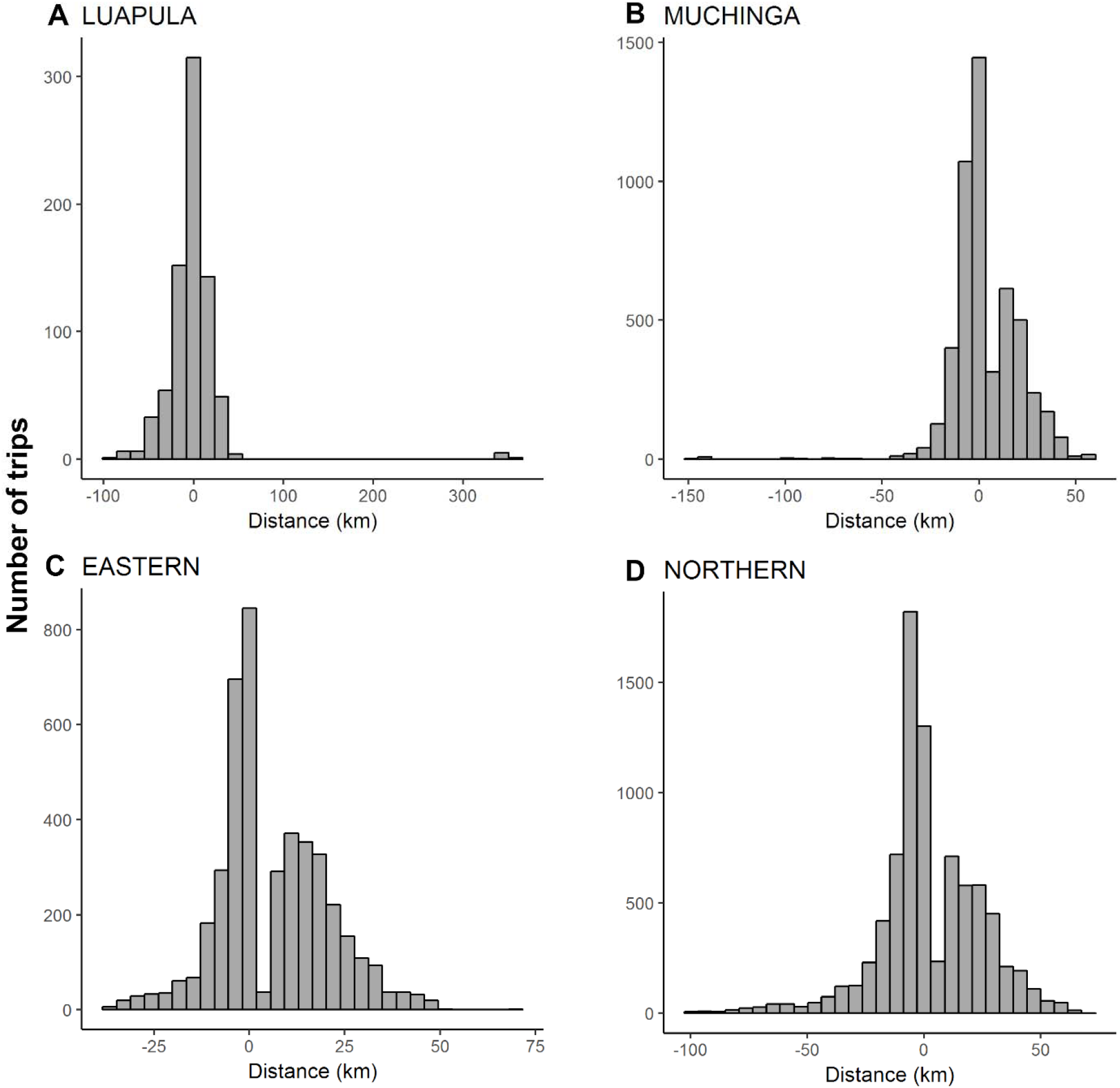
The range of differences in distance, for the four provinces (A-D) between Euclidean distance and optimal network route distance from city centers to nearest target community.

### Within district prioritization

In the first strategy examined, district-by-district community ranked prioritization strategy occurs, allocating spray to the top 50% risk households based upon use of the Malaria Atlas Project (MAP) risk approaches. As seen in Table 2, few communities were excluded from any prioritization strategy (4.5%) and few were included within all prioritization strategies (4.9%), indicating low levels of agreement among MAP risk strategies. In pairwise strategy comparisons, agreement was statistically better than chance between *Pf*PR and *An. funestus* risk, but no agreement between other prioritization strategies was observed (Additional Table 2). The overall Kappa statistic of agreement between the different prioritization strategies was −0.039.

**Table 2.** Number of communities to receive IRS, under the four risk prioritization strategies (*Pf*PR, *An. arabiensis, An. funestus, An. gambiae*) when spraying half of all households *within each* district.

|  | Number of Communities | Percent of Communities |
| --- | --- | --- |
| Prioritized by zero strategies | 216 | 4.5% |
| Prioritized by one strategy | 1,150 | 24.1% |
| Prioritized by two strategies | 2,031 | 42.6% |
| Prioritized by three strategies | 1,139 | 23.9% |
| Prioritized by four strategies | 232 | 4.9% |

The distance to communities from a city center to deliver IRS ranged from 53m to 115km, with 75% of communities being located with 25km of a city center. Differences in distances to reach communities were higher in prioritized communities compared to non-prioritized communities for *Pf*PR and *An. funestus* risk, and distances were lower in prioritized communities when based on *An. gambiae* risk (Table 3).

**Table 3.**
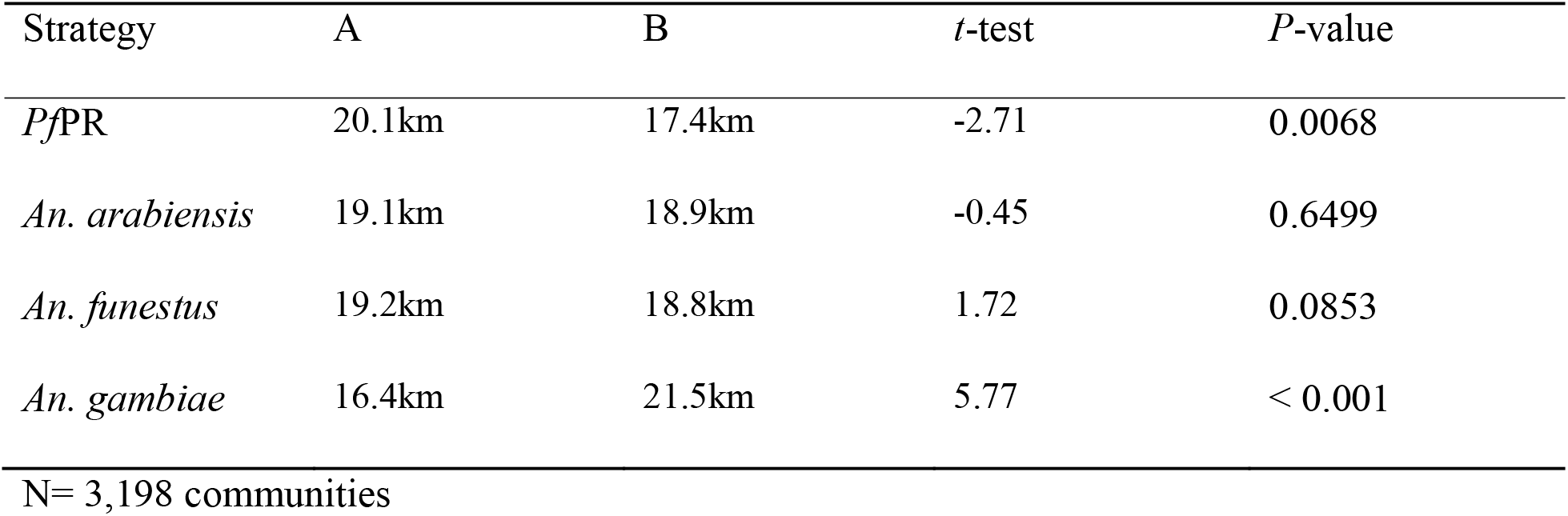
Mean distance to communities from city center for prioritized (A) and non-prioritized (B) communities by different strategies, *within each* district. (*t*-tests on log-transformed values).

| Strategy | A | B | *t*-test | *P*-value |
| --- | --- | --- | --- | --- |
| *PfPR* | 20.1km | 17.4km | -2.71 | 0.0068 |
| *An. arabiensis* | 19.1km | 18.9km | -0.45 | 0.6499 |
| *An. funestus* | 19.2km | 18.8km | 1.72 | 0.0853 |
| *An. gambiae* | 16.4km | 21.5km | 5.77 | < 0.001 |

N= 3,198 communities

### A cross-province prioritization

When considering deploying IRS across the entire four-province study area, using the same MAP prioritization approach, large variations are seen in the number of communities included by strategy, within province (Table 4), leading to high district-level variation in the allocation of IRS by strategy (Figure 3).

**Table 4.**
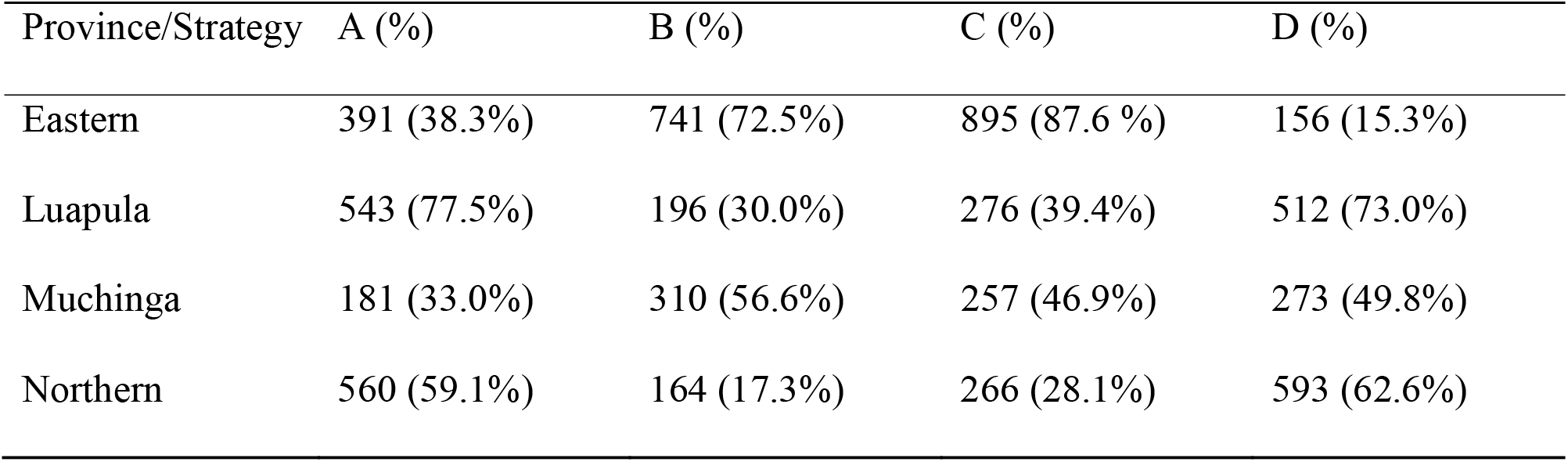
Number of communities (and percent) included for IRS by prioritization strategy (A. *Pf*PR, B. *An. arabiensis*, C. *An. funestus*, D. *An. gambiae*) when spraying half of all households *across all four* provinces.

**Figure 3.**
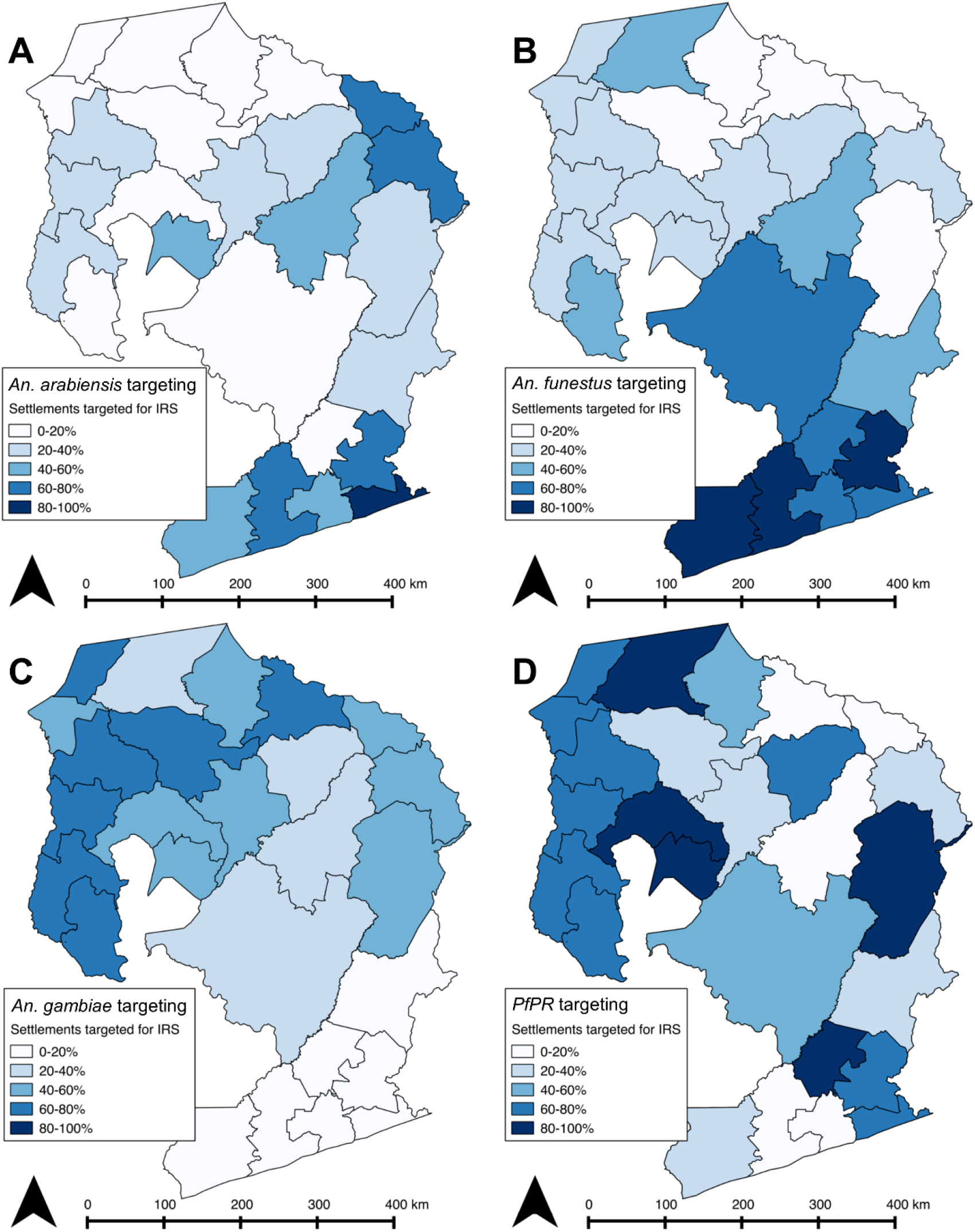
Percent of houses receiving IRS in each district by prioritization strategy (A. *Pf*PR, B. *An. arabiensis*, C. *An. funestus*, D. *An. gambiae*) when spraying half of all households *across all four provinces*.

Similarly to the district-by-district approach, there was large disagreement in communities covered between the different risk prioritization strategies. As seen in Additional Table 3, very few communities were excluded from any prioritization strategy (3.1%) and even fewer were included in all the prioritization strategies (2.6%). Comparing pairwise strategy sets, agreement was statistically better than chance between *Pf*PR and *An. funestus, PfPR* and *An. gambiae*, and *An. arabiensis* and *An. funestus*. The overall Kappa statistic of agreement between the different prioritization strategies was −0.103 (Additional Table 4).

The optimal route distance to communities ranged from 53m to 116km, with 75% of communities 25km or less from a city center. Distance to communities were no different between prioritized and non-prioritized communities for *An. gambiae* and *Pf*PR prioritization, while prioritized communities were statistically nearer to city centers based on *An. arabiensis and An. funestus* risk (Table 5).

**Table 5.** Mean distance to spraying communities from city headquarters for targeted (A) and untargeted (B) communities by different strategies, *across all four provinces*. (*t*-test on log-transformed values)

| Strategy | A | B | t-test | P-value |
| --- | --- | --- | --- | --- |
| *PfPR* | 19.7km | 18.1km | 1.51 | 0.1299 |
| *An. arabiensis* | 16.6km | 21.1km | 3.62 | <0.001 |
| *An. funestus* | 17.3km | 21.2km | 3.44 | <0.001 |
| *An. gambiae* | 19.3km | 18.7km | 0.56 | 0.5722 |

N= 3,198 communities

## Discussion

In this paper, models of allocating IRS were examined, using a combination of optimal network distributions based on road network routing, and spatial prioritization using risk maps estimating under 5 *Pf*PR, and suitability for the three major Anopheline vectors implicated in malaria transmission in Zambia. Differences were observed in the cost to prioritize communities with higher estimates of *Pf*PR and *Anopheles* species’ vector capacity, whether strategizing IRS application district-by-district, or across the four provinces in the study. In many cases, reaching communities prioritized by risk strategies did not differ significantly in cost from reaching non-prioritized communities, and were in some cases cheaper. Prioritizing communities for intervention at the provincial level rather than equally allocating coverage across districts led to variation in the proportion of houses receiving IRS at the district level, as shown in Figure 3. The findings of this study suggest areas with higher malaria burdens may not necessarily be more remote than with lower malaria burdens.

Additionally, a complete lack of agreement in IRS allocation between *Pf*PR and vector prioritization strategies presents a challenge to malaria programs, requiring programs to pick which measure of risk is most appropriate given their context. Risk maps are not often used when planning malaria interventions [33], and there is little literature to suggest which prioritization strategy has the most impact on reducing malaria. Further, these analyses utilized global risk maps of malarial risk indicators rather than local risk maps. The use of global risk maps can be considered both a strength and a limitation, with data availability being one of the primary strengths. Two of the most recent malaria risk maps in Zambia were subnational, and even sub-provincial, and so would not be useful for national IRS campaigns [34,35]. These risk maps need to be validated on a larger scale before they can be useful for malaria control programs. It remains to be seen whether localized, more specific risk maps have better agreement in prioritizing communities to receive limited IRS resources.

It has been noted by several authors that availability of accurately mapped road networks greatly limits the application of optimizing network routing models in the developing world [19,20]. We demonstrated the differences in estimating route costs between using simple Euclidian distance mapping, and network routing, suggesting this is an important gap to fill for effective planning for distribution programs of many kinds. In this study, the best available data were used, but results must be interpreted with the data limitations in mind. Forty percent of the communities enumerated were not captured by the network analysis because they were not within 5,000 meters of the road network and were therefore removed from further analyses. It is likely that the road network data is incomplete, rather than that these communities are indeed more remote than their counterparts nearer to the available road data. Indeed, there was no difference in malaria risk between captured and non-captured communities by any of the four measures used herein. Investment into geospatial data such as road networks would improve predictive modeling and precision public health delivery [36] of interventions such as IRS.

Further, while we believe we have developed a robust model there is always notable variability in IRS operations; for example, team size and number and location of operational bases may differ across a country context, and even within a province or district. It may be that the degree of this variability differs in harder to reach areas. While this model does not fully capture that variability, we believe it still offers a basis of comparison across the prioritization strategies and creates a framework for adding such complexity in the future. We note that these are findings from one country, Zambia, and that while the methods may prove practical to assess

IRS distribution in other countries, the results may differ due to differing population density and distribution, as well as transmission patterns.

## Conclusions

Areas with greater malaria burden or risk of malaria transmission are not necessarily more costly to reach for intervention delivery. A lack of agreement between different risk maps may be challenging for malaria control programs deciding how to prioritize where to spend resources.

## List of abbreviations

IRS: indoor residual spraying
ITN: insecticide-treated mosquito net

## Declarations

Ethics approval and consent to participate

Not applicable

## Consent for publication

Both the United States President’s Malaria Initiative and the Zambian National Health Research Authority reviewed the article before approving submission to a scientific journal.

## Availability of data and material

The satellite enumerations used in these analyses are proprietary data owned by the Zambian Ministry of Health, and request to access these enumerations can be made to the Zambian Ministry of Health. All other data used in these analyses are publicly available.

## Authors’ contributions

SJR and DL conceived of the analytical framework; SJR, DL, and BW conducted analyses; SJR, DL, BW, AM, AW wrote and edited the manuscript. All authors approved the final version of the manuscript.

## Acknowledgements

Not applicable

## Additional File

**Additional Table 1.** Number of city center points and communities, and number included in network analyses, for the four study provinces in Zambia

|  | Luapula | Muchinga | Northern | Eastern | Total |
| --- | --- | --- | --- | --- | --- |
| City center points | 50 | 66 | 59 | 61 | 236 |
| *City center points captured* | *50* | *61* | *58* | *59* | *228* |
| Total Communities | 758 | 5088 | 8241 | 4350 | 18,437 |
| *Communities captured* | *693* | *3140* | *5050* | *2363* | *11,146* |

**Additional Table 2.**
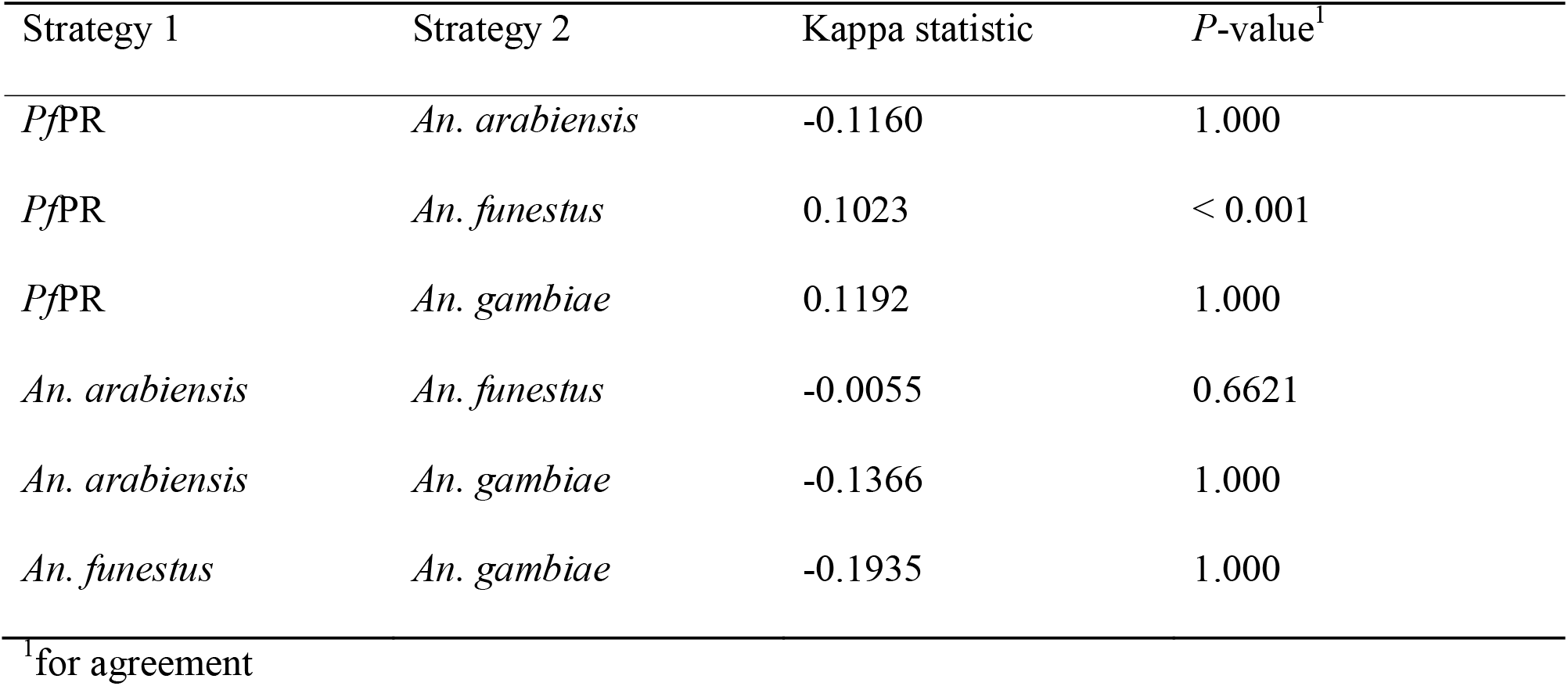
Kappa statistics for prioritizing communities (N=3,218) between pairs of prioritization strategies, *within each* district.

**Additional Table 3.** Number of communities to receive IRS, as prioritized under the four strategies (*Pf*PR, *An. arabiensis, An. funestus, An. gambiae*) when spraying half of all households *across all four provinces*.

|  | Number of communities | Percent of communities |
| --- | --- | --- |
| Prioritized by zero strategies | 148 | 3.1% |
| Prioritized by one strategy | 1,186 | 24.9% |
| Prioritized by two strategies | 2,296 | 48.2% |
| Prioritized by three strategies | 1,012 | 21.2% |
| Prioritized by four strategies | 126 | 2.6% |

**Additional Table 4.** Kappa statistic for prioritizing communities (N=3,218) between pairs of prioritization strategies, when spraying half of all houses *across all four provinces*.

| Strategy 1 | Strategy 2 | Kappa statistic | *P*-value<sup>1</sup> |
| --- | --- | --- | --- |
| *PfPR* | *An. arabiensis* | -0.2483 | 1.000 |
| *PfPR* | *An. funestus* | 0.0261 | 0.0359 |
| *PfPR* | *An. gambiae* | 0.1127 | < 0.001 |
| *An. arabiensis* | *An. funestus* | 0.2332 | < 0.001 |
| *An. arabiensis* | *An. gambiae* | -0.2812 | 1.000 |
| *An. funestus* | *An. gambiae* | -0.4595 | 1.000 |

<sup>1</sup>for agreement

